# Cancer drug sensitivity prediction from routine histology images

**DOI:** 10.1101/2023.06.03.543536

**Authors:** Muhammad Dawood, Quoc Dang Vu, Lawrence S. Young, Kim Branson, Louise Jones, Nasir Rajpoot, Fayyaz ul Amir Afsar Minhas

## Abstract

Drug sensitivity prediction models can aid in personalising cancer therapy, biomarker discovery, and drug design. Such models require survival data from randomized controlled trials which can be time consuming and expensive. In this proof-of-concept study, we demonstrate for the first time that deep learning can link histological patterns in whole slide images (WSIs) of Haematoxylin & Eosin (H&E) stained breast cancer sections with drug sensitivities inferred from cell lines. We employ patient-wise drug sensitivities imputed from gene expression based mapping of drug effects on cancer cell lines to train a deep learning model that predicts sensitivity to multiple drugs from WSIs. We show that it is possible to use routine WSIs to predict the drug sensitivity profile of a cancer patient for a number of approved and experimental drugs. We also show that the proposed approach can identify cellular and histological patterns associated with drug sensitivity profiles of cancer patients.

**Highlights:**

- Predicting drug sensitivity from routine histology images and cell lines
- Discovery of histology image patterns linked to drug sensitivity
- A novel deep learning pipeline for analysing drug sensitivity profiles

## 1 Introduction

The premise of precision medicine is to develop therapies that target important characteristics (such as the molecular profile) of an individual tumour. The outcome of drug therapy is often unpredictable (ranging from desirable to toxic) and predominantly driven by a tumour’s molecular profile [1]. The response to anti-cancer drugs can be influenced by both germline and acquired somatic mutations [2] as well as the status of molecular/signalling pathways [3], suggesting that therapies targeting genomic landscape of an individual are more effective compared to one-size-fits-all therapy approaches [4]. Pharmacogenomics is a pivotal component of precision oncology that fuses pharmacology and genomics to study an individual’s response to drug based on their genomic profile [5]. Recent advances in high-throughput drug screening and the availability of pharmacological data together with a multitude of omics data (genomic, mutational, transcriptomic, proteomic and metabolomic data) has paved the way for identifying genetic biomarkers that are associated with treatment response [6], [7].

Cancer cell lines (CCLs) provide an easy-to-manipulate vehicle for high-throughput drug screening at scale, prior to the more expensive *in vivo* testing and clinical trials of a drug [6]. Pioneers of these large scale genomic and drug screening datasets include the NCI-60 database [8], Cancer Cell Line Encyclopedia (CCLE) [9], Genomics of Drug Sensitivity in Cancer (GDSC) [10], and the Cancer Therapeutic Response Portal (CTRP) [11]. These datasets have helped assess the sensitivity of many compounds including FDA approved drugs *in vitro* and have led to the discovery of novel anti-cancer therapies [12], [13]. The pharmacogenomics data provided by these initiatives have enabled collective analysis of drug sensitivity and gene expression data to uncover novel drug-gene relationships [8]. Several machine learning (ML) methods have been proposed for associating mutation and gene expression data of cancer cell lines with their respective drug efficacy metrics such as half maximal inhibitory concentration (IC_50_), or area under the dose response curve (AU-DRC) [10], [14], [15]. Similarly, deep learning (DL) based methods [16]–[19] and several other approaches [20]–[22] have been proposed for predicting patient response to drugs using genomic information.

Despite advances in genomics-based drug sensitivity analysis, the applicability of genomics profiling for selecting appropriate drugs remains limited. The digitisation of tissue slides and recent advances in detailed tissue profiling using digital scans of routine H&E tissue slides offers a novel way to predict drug sensitivity via spatial histological profiling. To the best of our knowledge, this is the first study that proposes prediction of sensitivity to multiple drugs from routine H&E images by training a predictive model using CCLs imputed drug sensitivity data. We address the question as to whether and to what extent it is possible to predict a breast cancer patient tumour’s sensitivity to multiple approved and experimental drugs on the basis of their histological profile as captured by DL based analysis of the H&E images. The resulting association of visual histological patterns with drug sensitivity can be helpful in identifying histological motifs associated with high and low sensitivity of drugs. Not only can it pave the way for spatial characterisation of treatment response, it also carries the potential of ruling out treating a patient with certain drugs due to their histological profile.

Histological examination of tissue sections is considered a gold standard for the clinical diagnosis of solid tumours. Recent advancements in deep learning for computational pathology have proven valuable for using WSIs of routine Haematoxylin and Eosin (H&E) stained tissue sections to predict cancer subtypes [23], [24], patient survival [25], [26], mitosis detection [27], DNA methylation patterns [28], cellular composition [29], [30], and tumour mutation burden [31]. Moreover, histology image based prediction of mutation and expression profile of different genes [32]–[34] and prediction of molecular markers and pathways [35], [36], has been achieved using deep learning. Recently, a deep learning model has been proposed for predicting breast cancer patients gene expression state from WSIs [37].

In this work, we investigate the association between cellular and morphometric patterns contained in the digitised WSIs of routine H&E tissue slides of breast cancer tumours and their drug sensitivity profiles. We employ the method proposed in [37] for predicting patient’s likelihood of response to treatment to specific drugs. The framework proposed in this study (see **Fig 1**) offers several possible advantages. First, it enables direct association of phenotypic information present in WSIs with the likelihood of response to different drugs. Second, by utilising pharmacogenomic datasets, a large population of patients can be virtually screened for a broad spectrum of compounds in a relatively short amount of time giving valuable insights into the association of patient-specific histological signatures with drug sensitivities. Third, harnessing ML for drug sensitivity estimates allow modelling the relative contribution of expression level of various genes to the patient’s sensitivity to a broad spectrum of compounds in an unbiased manner. The proposed framework does not rely on any assumption about the mechanism of action of compounds, which may be unknown, or on patient’s survival data. Fourth, the framework offers flexibility to investigators when studying the response of a lead compound in different tissue types, subtypes, or any other patient population as part of a drug discovery pipeline. Finally, histological patterns discovered using the proposed framework can be easily translated into clinical practice as it is built on digital scans of routine H&E tissue slides and does not require any expensive or time-consuming assays.

**Figure 1:**
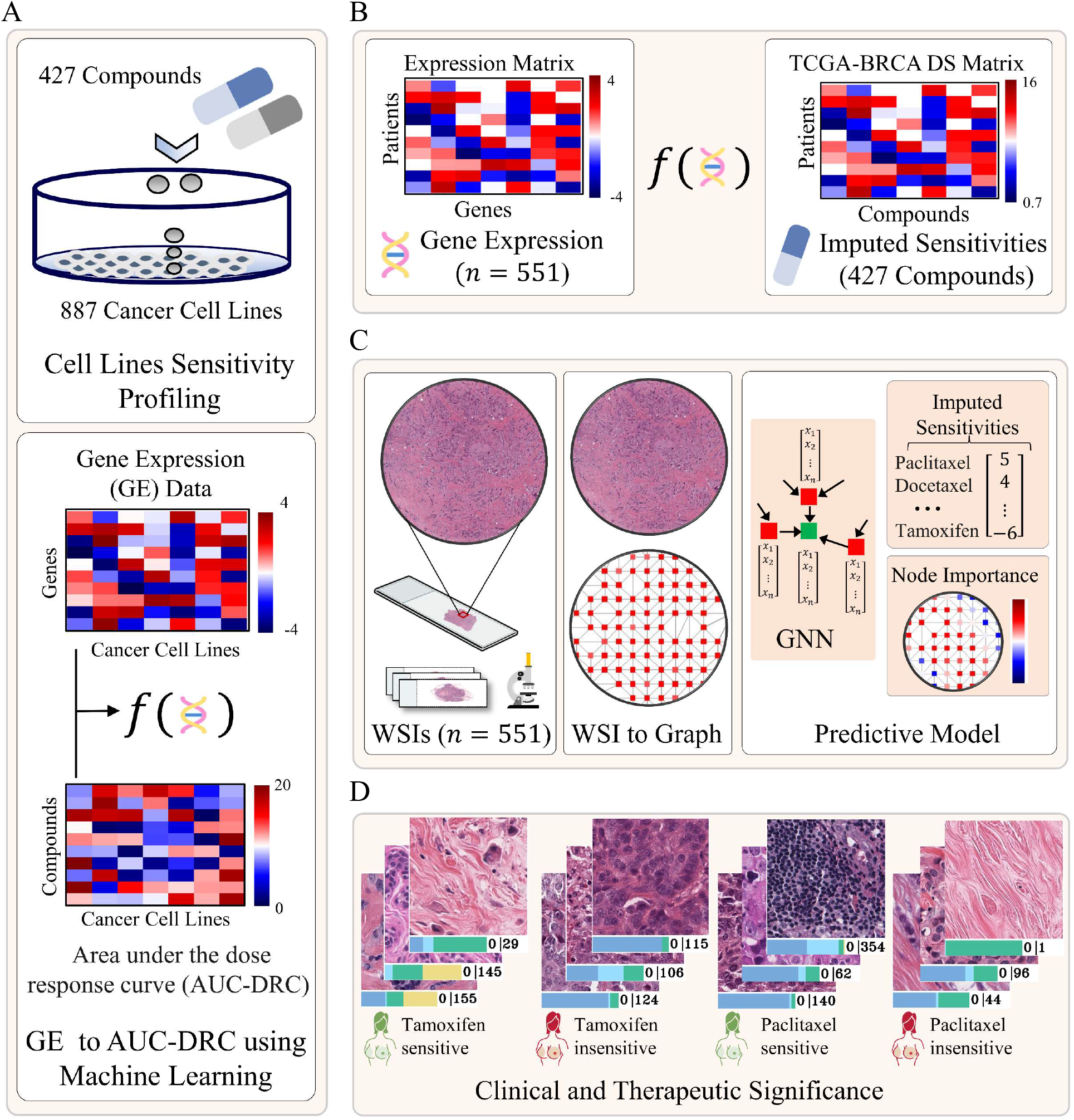
Workflow of the proposed approach for predicting patient sensitivities to different drugs from histology images. A) A regression model was developed using cancer cell line gene expression data and *in vitro* drug screening data to learn association between cell lines gene expression profile and their sensitivity to 427 compounds. B) The trained model was then used to infer the sensitivity of breast cancer patients to these drugs based on their gene expression. The output of the model is a matrix listing the gene expression based imputed drug sensitivities of each patient (one per row) to 427 compounds (one per column). C) Prediction of patient sensitivities to compounds from whole slide images (WSIs) of formalin-fixed paraffin-embedded (FFPE) H&E stained tissue section using a Graph Neural Network (GNN). We represent each WSI as graph and then pass the WSI-graph as input to a GNN for predicting WSI-level and patch-level sensitivities of patient to different drugs. Node-level prediction highlight the spatially resolved contribution of different region of WSI towards the predicted sensitivity of a certain drug. D) Histological motifs associated with high and low sensitivity of Tamoxifen and Paclitaxel are shown for illustration

## 2 Results

### 2.1 Imputed drug sensitivities from cell lines

We used estimated sensitivities of TCGA-BRCA patients (*n*=551) to 427 compounds from a previously published method [15] which fits a linear ridge regression model between CCL based gene expression data of genes as input and the corresponding *in vitro* measurement of drug response in terms of area under the dose response curve (AU-DRC) as output. Once trained, these linear regression models (one per drug) were then used for imputing the sensitivity of the Cancer Genome Atlas breast cancer (TCGA-BRCA) patients to every drug in the CTRP database [11]. An overview of approach used for obtaining patients ground truth sensitivity estimates is provided in **Fig 1A** and **Fig 1B**. It is important to note that the higher the AU-DRC, the lower the drug sensitivity since a higher concentration of the drug is required for it to be effective and vice versa.

### 2.2 Analytical Pipeline for whole slide image analysis and predictive modelling

To explore the association between cellular and histological patterns contained in WSIs and patient tumour’s sensitivity to different drugs, we propose an end-to-end DL pipeline that takes WSI of a patient as input and predicts the sensitivity of 427 compounds as output. An overview of the proposed framework is provided in **Fig 1C**. We employed our in-house *SlideGraph*^*∞*^pipeline [37] that first constructs a graph representation of the WSI and then uses a graph neural network (GNN) to predict node-level (patch-level) and WSI-level sensitivity of a patient to all the compounds (compounds listed in Table S1). The node-level scores are then used to identify regions within the WSI that contribute to high or low sensitivity. This gives insight into cellular composition and diversity in terms of different types of cell present in the tumour microenvironment (TME) in a spatially resolved manner as shown in **Fig 1C** and **1D**. In this study, we utilised WSIs of TCGA-BRCA patients (*n* = 551) that have gene expression based imputed sensitivity score for all 427 drugs.

### 2.3 Prediction of drug sensitivity from whole slide images

Our predictive analysis shows that patient sensitivity to a wide range of compounds can be predicted from histology images with high Spearman Correlation Coefficient (SCC) values and significant *p-*values, as shown in **Fig 2A**. From the plot, it can be observed that for 186 out of 427 drugs, sensitivities predicted by our model are statistically significantly correlated (*p* ≪ 0.001) with the ground-truth sensitivity estimates. **Fig 2A** also shows the top 10 drugs that are predicted with high SCC values are shown as diamonds, with each drug represented by a unique colour. The distribution of SCC for the highlighted drugs is shown in **Fig 2B**. From the boxplot, it can be observed that all the listed drugs have their mean SCC values greater than 0.5. This shows that the responsiveness of the patient’s tumour to a broad spectrum of drugs can be inferred from their histological imaging profile.

**Figure 2:**
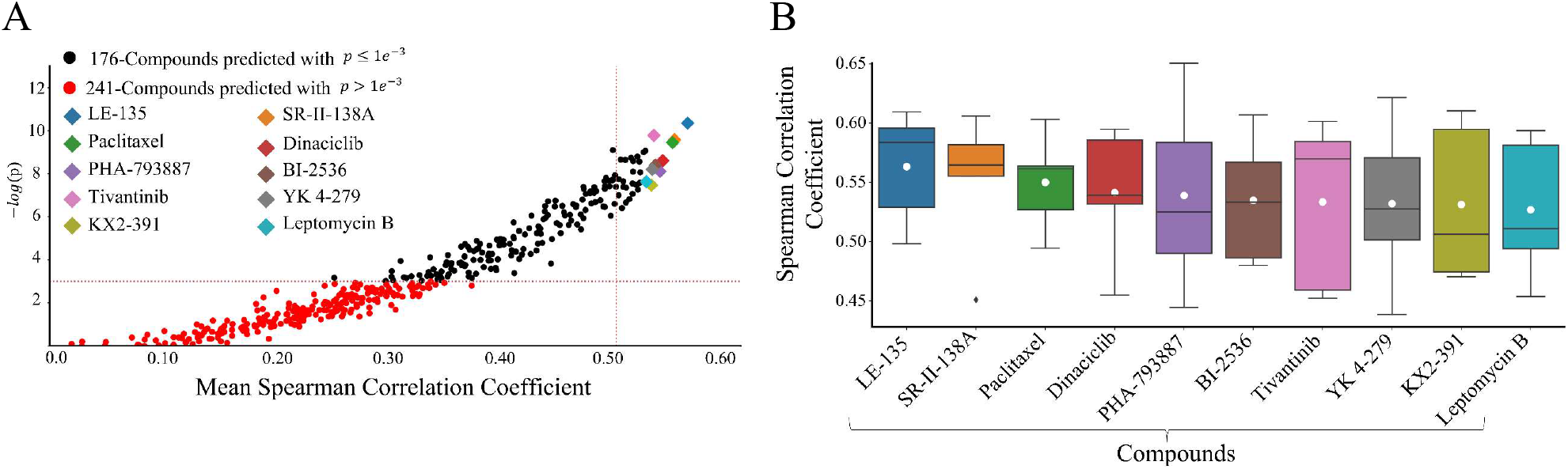
Predictibility of patient tumour sensitivity to different compounds from histology images. A) Scatter plot showing the correaltion between histology-image based predicted sensitivity and gene expression based imputed drug sensitivity using spearman correlation coefficent (SCC) as performance metric. The mean SCC across 5-cross validation folds is shown along x-axis with its corresponding -log10 (FDR corrected p-value) along y-axis. Each dot represents a particular drug and its colour represents the statistical significance of the alignment between the predicted score and ground truth value. The diamond highlight the top-10 drugs whose histology image based predicted sensitivity match significantly with ground truth values in terms of SCC. B) Boxplot showing the distribution of SCC across 5-cross validation for the top-10 best predicted compounds from histology images. The white dot within each boxplot highlight the mean SCC across different runs.

### 2.4 Association of drug sensitivity with spatially resolved cellular and histopathological phenotypes

Pathological assessment of tumour microenvironment (TME) in breast cancer plays a pivotal role in predicting tumour behaviour and treatment outcome [38]. The proposed graph-based approach allows highlighting spatially localised morphometric patterns associated with the sensitivity of different drugs using node-level prediction score as a guiding signal. **Fig 3** shows some example heatmaps highlighting the contribution of different regions of the WSI towards the predicted sensitivity of the patient’s tumour to paclitaxel and tamoxifen. For both drugs, an example WSI and a heatmap are shown for a highly sensitive tumour and a relatively insensitive tumour. The heatmaps show the relative contribution of different regions of the WSI toward the predicted sensitivity estimate of the tumour to a certain drug using pseudo-colours, with dark red colour indicating regions contributing to high sensitivity and dark blue colours corresponding to regions contribution to the prediction of low sensitivity. From the high and low contributing regions, we extracted some sample regions of interest (ROIs) outlined by red and blue colour, respectively, in **Fig 3**. The figure illustrates that ROIs associated with high sensitivity to paclitaxel exhibit relatively high proportions of tumour cells as well as lymphocytes. This aligns with previous research demonstrating that tumour infiltrating lymphocytes (TILs) can act as independent predictor of a patient tumour’s sensitivity to chemotherapy drugs [39]. Conversely, ROIs indicative of low sensitivity to paclitaxel are characterized by a notable myxoid change in the stroma. This feature has previously been associated with poor overall survival (OS) and relapse-free survival (RFS) in triple-negative breast cancer [40].

**Figure 3:**
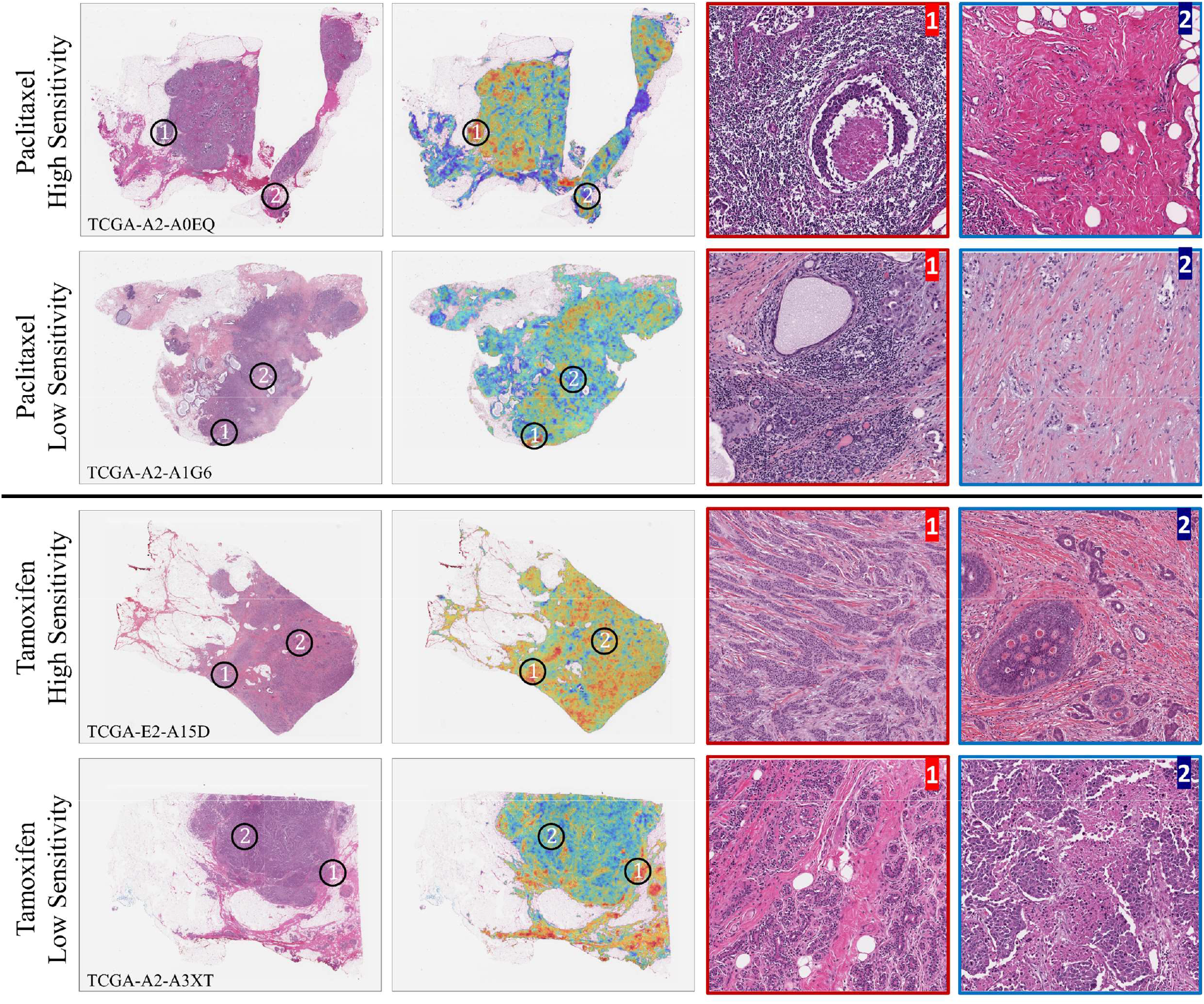
Illustration of different histological patterns within the WSIs associated with patient sensitivity toward Paclitaxel and Tamoxifen. Example WSIs and their corresponding heatmaps are shown for patients being either highly or lowly sensitive to these drugs. The heatmaps use pseudo colours (blue to red) to highlight the spatially resolved contibution of different regions of the WSI toward the predicted sensitivity. Bluer and redder colour respectively indicate regions of the WSI that contribute the most toward deciding low or high sensitivity. From the WSIs, we extracted magnified versions of regions of interest (ROIs), indicated by the black circles within the WSIs, that are associated with high and low sensitivity of a certain drug. ROIs outlined in red colour are indicative of high sensitivity, while those outlined in red blue colour are indicative of low sensitivity of a certain drug.

Regarding tamoxifen, ROIs indicative of high sensitivity are characterised by tumour cells that have relatively low nuclear pleomorphism. Conversely, in ROIs indicative of low sensitivity, the presence of necrosis, increase in mitotic count and cribriform DCIS (ductal carcinoma in-situ) can be observed. This is in line with previous studies that have found correlation of necrosis with larger tumour size and higher cancer grade, and hence it may be postulated that highly necrotic regions may contribute to relatively low sensitivity to tamoxifen [41].

### 2.5 Histological patterns associated with sensitivity to different drugs

We investigated association of visual histological patterns in the WSIs with the sensitivity of drugs by identifying exemplar patches (of size 512 × 512 pixels at a spatial resolution of 0.25 microns-per-pixel) for the high and low sensitivity of each drug using clustering. For these patches, we also computed the cellular composition (counts of neoplastic, inflammatory, connective, and epithelial cells), overall cellularity and mitotic counts. These visual patterns or histological motifs can be used as a potential indicator to guide therapeutic decision making. **Fig 4** shows representative patches for patients showing high or low sensitivity to paclitaxel and tamoxifen. The most prominent feature in patches representative of high sensitivity to paclitaxel is sheets of pleomorphic tumour cells. Additionally, some patches also exhibit evidence of necrosis and lymphocytic infiltration. In contrast, dense sclerotic stroma is the most consistently observed feature across patches representative of low sensitivity to paclitaxel. Regarding tamoxifen, we observed similar histological patterns relating to high sensitivity as the ones present in patches predicting low sensitivity to paclitaxel. For example, both tamoxifen high sensitivity and paclitaxel low sensitivity representative patches are more sclerotic with less pleomorphic tumour cells. However, in patches indicative of patient low sensitivity to tamoxifen the cells are more pleomorphic with some evidence of necrosis.

**Figure 4:**
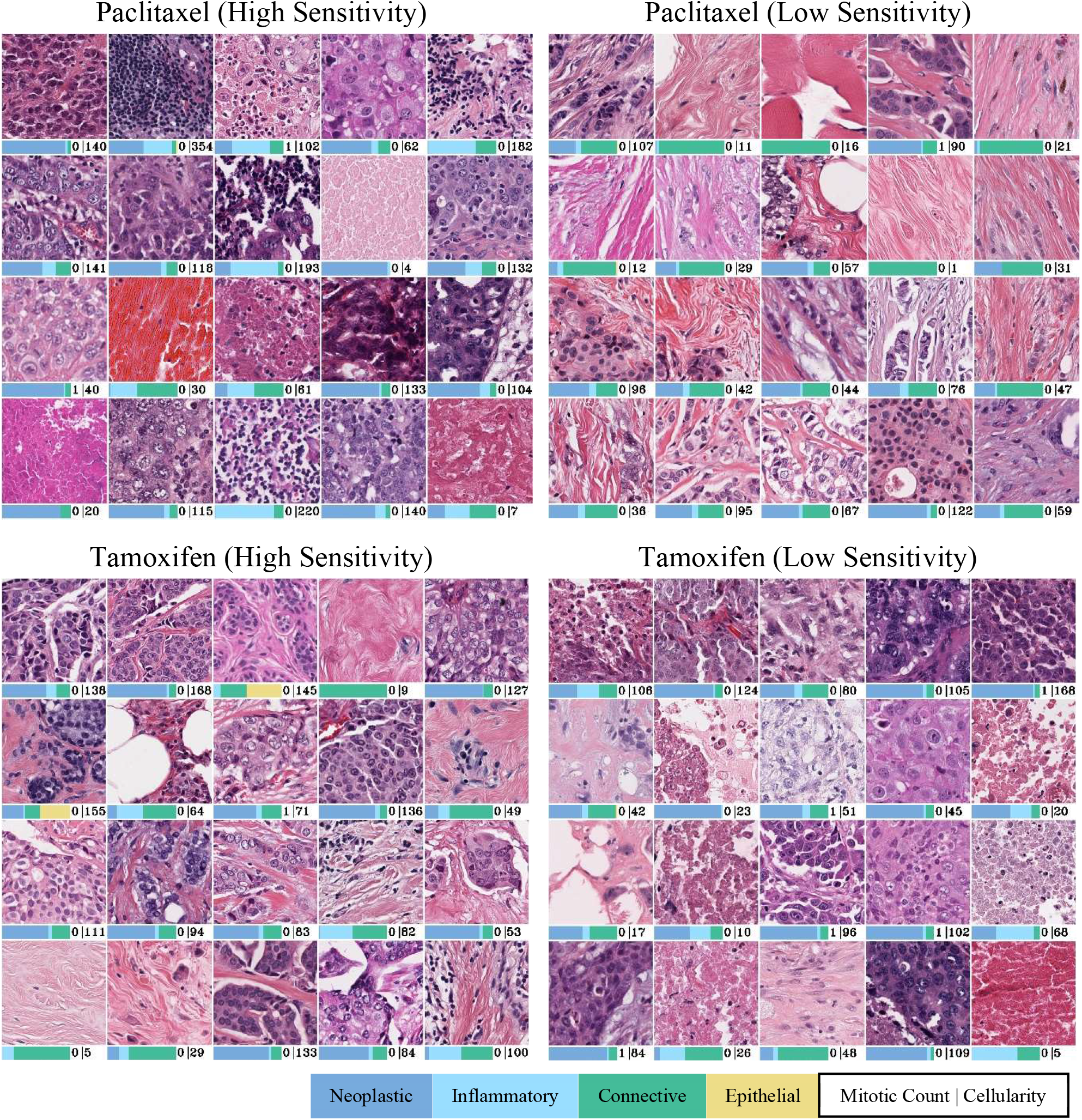
Representative patches of Paclitaxel and Tamoxifen high and low sensitivity. The bars below each patch respectively show its cellular composition in terms of relative counts of four different cell types (neoplastic, inflammatory, connective and epithelial), mitotic counts and overall cellularity (cell counts). Sheet of pleomorphic tumor cells (e.g. row1 image 1 and 4; row 2 image 1, 2 and 5; row 3 image 1 and 5; row 4 image 2 and 4; row 5 images 1 and 5), lymphocytic infiltrate (row 1 image 2 and 5, row 4 image 3 and row 5 image 3) and necrosis (row 1 image 3, row 2 image 4, row 3 image 3, row 4 image 1, row 5 images 2 and 4) can be seen in patches relating to paclitaxel high sensitivity. In patches relating to paclitaxel low sensitivity and tamoxifen high sensitivity, the consistent feature is dense sclerotic stroma. In patches associated with tamoxifen low sensitivity, the cells are more pleomorphic (row 1 images 2 and 4, row 2 images 3 and 4, row 3 image 3) with some evidence of necrosis (row 1 image 1, row 2 image 5, row 3 images 2 and 5, row 4 image 2 and row 5 image 1).

### 2.6 Association of drug sensitivity with pathologist-assigned histological phenotypes

We validated the predicted sensitivity estimates for several drugs and their associated histological patterns (identified using the proposed pipeline) by calculating Kendall’s tau correlation between image-based predicted sensitivities of drugs and pathologist assigned WSI-level histological phenotypes. We show an example heatmap in **Fig 5** where strong correlations between sensitivity of several drugs and a number of histological phenotypes can be seen. Overall, the predicted sensitivities of almost all chemotherapy drugs listed in the figure (e.g., paclitaxel, docetaxel, doxorubicin, etc,) shows strong positive correlation with cancer grade, mitosis, inflammation, necrosis, nuclear pleomorphism, epithelial tubule formation, TIL regional fraction and buffa hypoxia score. The opposite is true for sensitivity to tamoxifen sensitivity, which is a hormone therapy drug, as can be seen in the figure. For example, patients predicted as less responsive to tamoxifen tend to exhibit low grade cancer with reduced hypoxic tumour features. On the other hand, an opposite pattern can be observed for paclitaxel, a chemotherapy drug, in terms of drug sensitivity and tumour characteristics.

**Figure 5:**
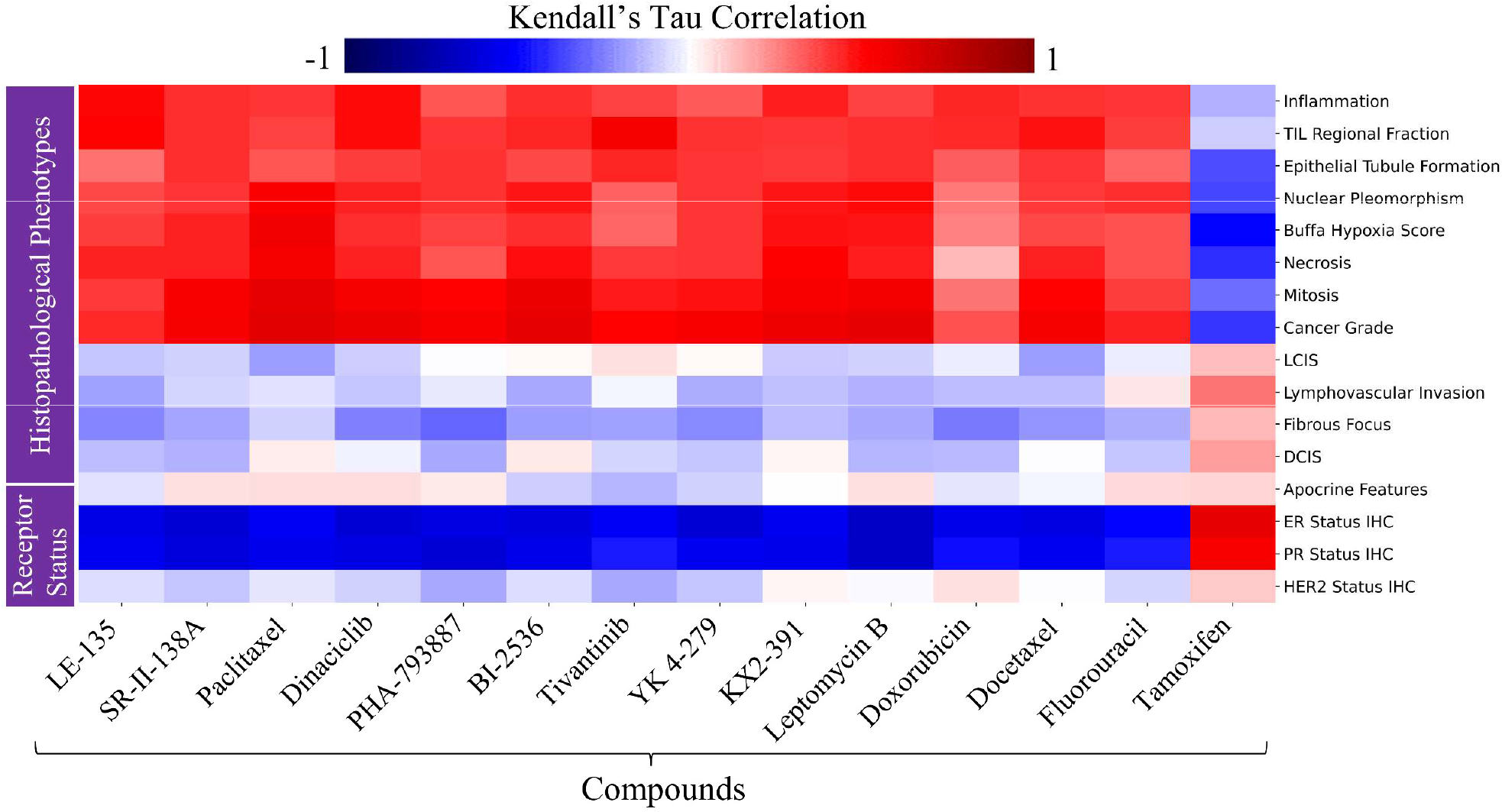
Association of drug sensitivity predicted by our model with pathologist assigned WSI-level histological phenotypes and breast cancer receptor status. Drugs are shown along x-axis, and histological phenotypes and receptor status are shown along y-axis. Red and blue colours indicate the degree of association between the predicted sensitivity of drugs and a specific histopathological phenotype or clinical marker. Bluer colour show strong negative correlation while strong positive correlation is shown using dark-red colour. (Abbreviation: TIL: Tumour Infiltrating Lymphocytes, LCIS: Lobular Carcinoma in situ, DCIS: Ductal Carcinoma in situ)

### 2.7 Association of drug sensitivity with receptor status

We found that patients sensitivities to drugs can be explained in terms of routine breast cancer clinical biomarker (ER, PR and Her2 status). **Fig 5** shows the association of image-based predicted sensitivities with ER, PR and Her2 status. From the figure, patient sensitivities to almost all listed chemotherapy drugs (e.g., paclitaxel, docetaxel, etc,) show negative association with ER and PR status positivity, whereas the opposite is true for tamoxifen. Apart from ER and PR status, slight correlation of Her2 status can also be seen with sensitivities of different drug as evident from the figure. These results are in line with previous studies that have reported better 5-year overall survival (OS) of low-grade ER-positive patients treated with tamoxifen [42].

### 2.8 Correlation of drug sensitivity with cellular composition

We analysed the relative proportion of neoplastic, inflammatory, connective and epithelial cells in WSI patches contributing to high and low sensitivity, as shown by radar plot of the various cellular counts in **Fig 6**. For all drugs other than tamoxifen, image patches representative of high sensitivity have relatively higher proportion of inflammatory cells than in image patches corresponding to the low sensitive group. Apart from inflammatory cells, the sensitivity of different compounds shows association with different patterns of cellular composition. For example, paclitaxel and KX2-391 show high sensitivity when the counts of neoplastic cells are relatively higher. As for the remaining drugs, their counts of neoplastic cells do not show significant difference in high and low sensitive group. Similarly, BI-2536, Dinaciclib, Paclitaxel, and Leptomycin B show high sensitivity when the count of normal epithelial cells is relatively low, while the remaining drugs show high sensitivity when epithelial cells are higher in number. Among the listed drugs, a different pattern is shown by tamoxifen, which is highly sensitive when the normal epithelial cell count is relatively high, whereas its sensitivity is low in tumour-rich regions (i.e., patches with relatively high neoplastic cell counts with almost no normal epithelial cells). Finally, for all drugs the count of connective cells is roughly the same between high and low sensitivity groups.

**Figure 6:**
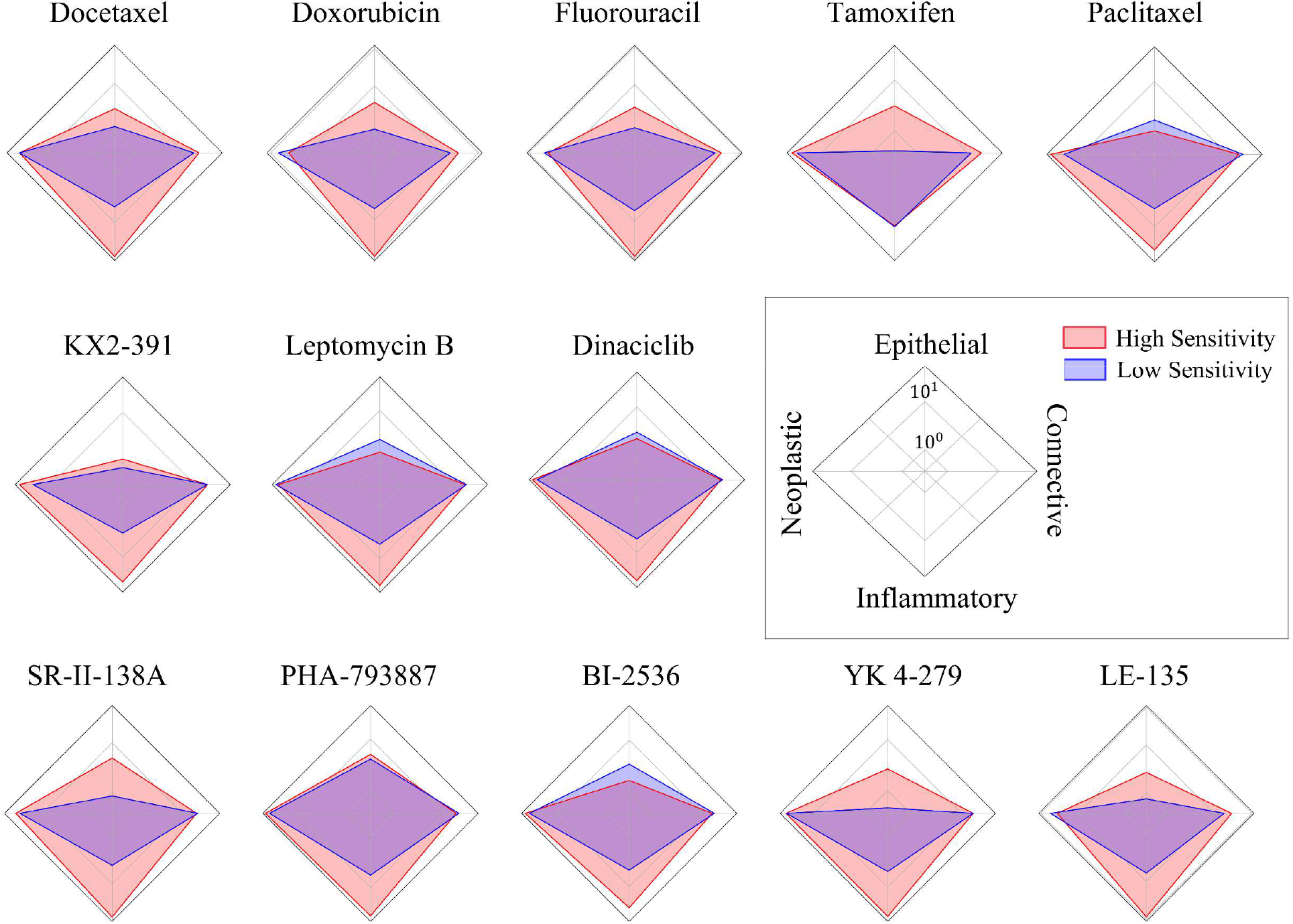
Radar plots of drug sensitivity and cellular composition. The plots show the association of relative counts of different types of cells with high and low sensitivity to a certain drug. Each axis of the plot represents a particular cell type, and the length of the axis shows their counts on a log scale. For example, the radar plot of Paclitaxel shows that patients who are highly sensitive to Paclitaxel have relatively higher numbers of inflammatory and neoplastic cells compared to those who are less sensitive.

### 2.9 Correlation between drug sensitivity and inflammatory to neoplastic cell ratio

We assessed the association between patch-level inflammatory to neoplastic cell counts ratio (INCCR) and sensitivities of several drugs. More specifically, for each drug we computed the patch-level INCRR in high sensitivity regions and plotted their distribution as boxplot, as shown in **Fig 7A**. From the figure, it can be observed that all chemotherapy drugs show high sensitivity when the patch-level INCCR is higher, whereas the opposite is true for tamoxifen. We analysed our findings statistically by computing the *p*-value between INCCR of high and low scoring patches of a certain drug using Wilcoxon rank-sum test. All drugs with * next to their name show statistically significant difference in INCCR ratio (*p* ≪ 0.05), while the ones without * (paclitaxel and tamoxifen) do not show statistically significant difference in INCCR between patches contributing to high and low sensitivity.

**Figure 7:**
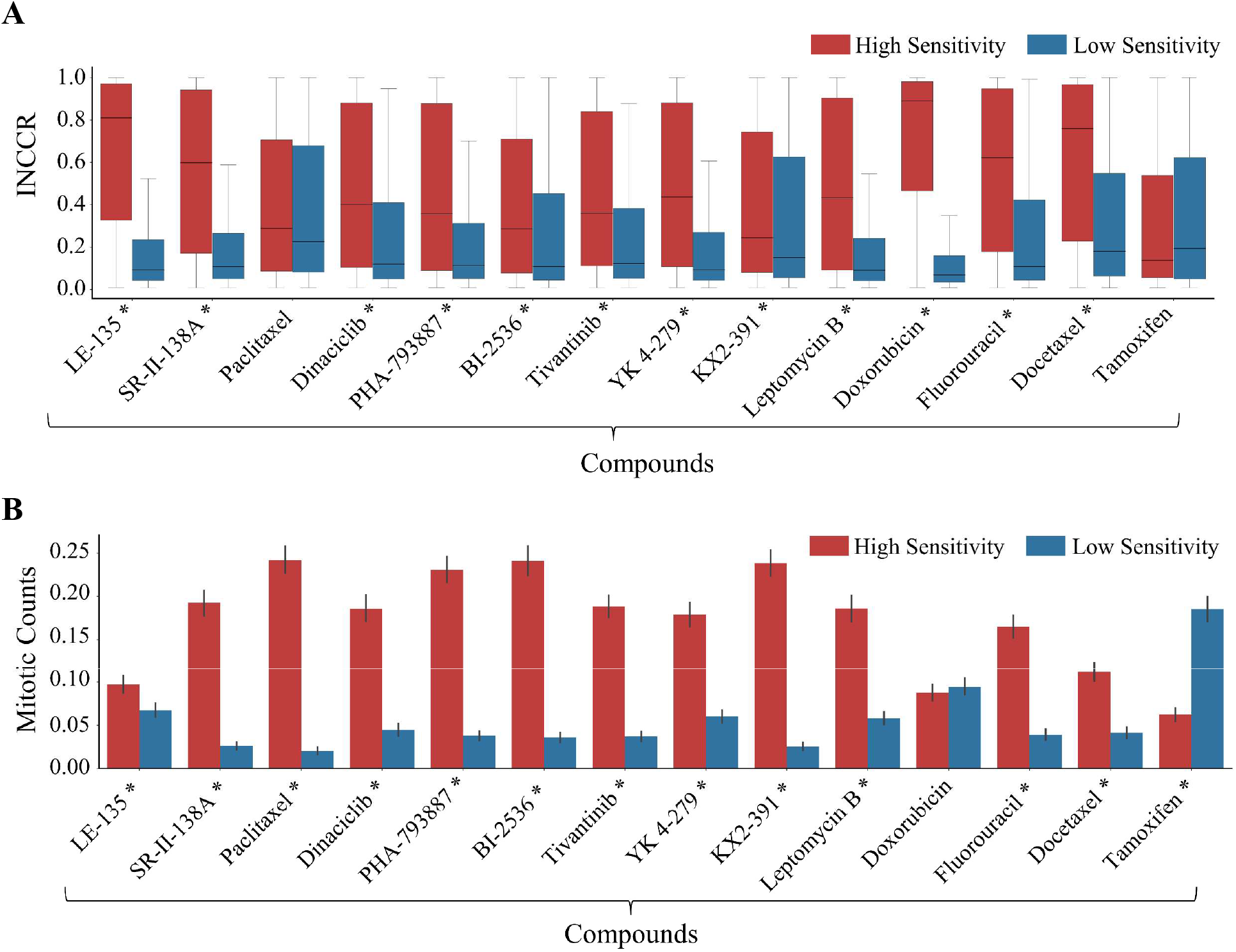
Statistical plots showing association of compounds sensitivity with; A) inflammatory to neoplastic cell counts ratio (INCCR) and B) mitotic counts. Compounds are shown along x-axis while the distribution of INCCR/Mitotic count is shown long y-axis. Red and blue colour represent the high and low sensitive group, respectively. Compounds with * next to their name show statistically significant difference in INCCR (in case of A) and mitotic counts (in case of B) between patches indicative of high and low sensitivity.

### 2.10 Correlation between drug sensitivity and patch-level mitotic counts

We found the sensitivities of drugs associated with patch-level mitotic counts. **Fig 7B** shows the distribution of patch-level mitotic counts in high scoring patches of a certain drug high and low sensitivity. From the figure, all the listed chemotherapy drugs (e.g. docetaxel, paclitaxel, doxorubicin, etc,) show high sensitivity when the patch-level mitotic counts are higher, whereas the opposite is true for tamoxifen which shows high sensitivity when patch-level mitotic counts are lower. We validated these findings statistically by computing the *p*-value between mitotic counts of high and low scoring patches of a certain drug using Wilcoxon rank-sum test. For all drugs listed in the figure we found statistically significant difference in patch-level mitotic counts (*p* ≪ 0.05) in high and low sensitivity patches.

## 3 Discussion

In this study, we proposed a novel approach to predict the sensitivity of breast cancer tumours to various drugs from histological patterns in their whole slide images (WSI). We employed our *SlideGraph*^*∞*^ pipeline [37] to explicitly model the local and global histological patterns in the WSI. More specially, we construct a graph representation of the WSI first and then train a graph neural network (GNN) that not only predicts WSI-level sensitivity but also highlights spatially resolved contribution of different WSI regions towards the predicted sensitivity estimate. We analysed the performance of *SlideGraph*^*∞*^ in predicting the sensitivity of TCGA breast cancer patients for 427 drugs when using only routine H&E histology images. Almost one half (186 out of 427 drugs) showed a statistically significant correlation between the patients’ predicted sensitivity to the drugs based on histology images and their imputed sensitivity based on gene expression. Moreover, we also identified histological patterns associated with high and low sensitivities to several drugs in terms of cellular composition, mitotic counts and histological motifs. Our analysis shows that patient sensitivity to a significant number of drugs can be predicted from histology images.

The proposed approach is fundamentally different from other approaches that predict patient sensitivity to different drugs by associating genomic and pharma data from cancer cell lines using machine learning. Most of these methods aim to discover patterns of gene expression that play a role in determining patient responsiveness to a particular drug. To the best of our knowledge, this is the first study to associate the sensitivity of anticancer drugs with tissue phenotypic information from routine histology images. We anticipate that direct association of drug sensitivity with histological profiles will help in discovering novel histological patterns that can be analysed by a pathologist to assist with therapeutic decision making for individual patients. The ground-truth sensitivity estimates used for training our histology-based model were obtained using a model trained on pharmacogenomic data of cancer cell lines. The motivation behind using cancer cell line for ground-truth sensitivity estimates is that it allows virtual screening of patients for a large number of compounds in a relatively short time and with minimal cost. We analysed the effectiveness of the proposed method on breast cancer data only, but it can be extended to other cancer types, subtype or patient population examined by the investigator. Finally, the proposed pipeline allows virtual screening of cellular response of biological specimen to lead compound studied during drug discovery.

Despite the promising results, the proposed approach has several limitations. As is the case with several other studies [43], the ground-truth sensitivity estimates used by our method are obtained based on gene expression data. While other studies [44], [45] have shown that drug sensitivity estimates based on gene expression profile are accurate and useful, extensive validation is still needed as gene expression may not be the only factor responsible for drug sensitivity. For example, epigenetic factors and proteomic expression changes can impact drug sensitivity without having any direct gene expression change. Another fundamental limitation stems from the use of ground-truth sensitivity estimates inferred from cancer cell lines which, while being low cost and high throughput, lack the microenvironment components that are known to influence response to therapy [46]. Although we performed extensive validation of sensitivities predicted by our model by association with manually assigned histological phenotypes, breast cancer receptor status and hypoxia scores, more stringent analysis is needed by validating the sensitivities predicted by our model using Randomized Control Trial (RCT) data as a possible future extension.

## Acknowledgments

The authors are grateful to Prof. Janes Armes (Sullivan Nicolaides Pathology, Sunshine Coast), Dr. Fouzia Siraj (Scientist D and Senior Pathologist at ICMR-National Institute of Pathology, India) and Emmanouil Karteris (Brunel University, London) for discussions and feedback on the topic of this study. MD would also like to acknowledge the PhD studentship support from GlaxoSmithKline and the Department of Computer Science at the University of Warwick. FM and NR were supported in part by the PathLAKE digital pathology consortium which is funded from the Data to Early Diagnosis and Precision Medicine strand of the government’s Industrial Strategy Challenge Fund, managed and delivered by UK Research and Innovation (UKRI). FM also acknowledges financial support from EPSRC EP/W02909X/1.

## Author Contributions

Original idea: FM and MD, Supervision: FM and NR, Funding acquisition: NR, KB and NR, Pathology review: LJ, Oncology input: LSY, Methods development: FM and MD, Results and Analysis: FM, MD and NR, Programming support: QDV. Writeup: All authors.

## 3.1 Methods

### 3.1.1 Dataset

#### 3.1.1.1 Acquisition of whole slide images and drug-response data

We collected 1133 Whole Slide Images (WSIs) of Formalin-Fixed Paraffin-Embedded (FFPE) Haematoxylin and Eosin (H&E) stained tissue section of 1084 breast cancer patients from the Cancer Genome Atlas (TCGA) [47], [48]. The gene expression profile based drug sensitivity estimates of 936 TCGA breast cancer patients for 427 compounds were obtained from the work of Gruener *et al*. [49]. To limit the impact of various artefacts, we excluded WSIs that met any of the following criteria: 1) containing extensive blurry areas; 2) having abnormal staining with minimal informative tissue regions; or 3) lacking baseline resolution information. After filtering, in total we used data for 551 patients along with their imputed sensitivity to 427 drugs in our analyses.

#### 3.1.1.2 Acquisition and filtering of whole slide images

## 3.2 Pre-processing

We first segment the tissue region of each WSI and exclude regions with tissue artifacts (pen-marking, tissue folding etc.,) using a tissue segmentation model. Each WSI is then tiled into patches of size 512 × 512 pixels at a spatial resolution of 0.25 microns-per-pixel (MPP). Patches with less than 40% of tissue region (pixels with intensity higher than 200) are discarded and the rest of patches (tumour and non-tumour) are used in the study. Target drug sensitivity data is converted into z-score prior to prediction with high sensitivity corresponding to lower area under the dose response curve for that compound and vice versa.

## 3.3 Graph Modelling of Whole Slide Image

A graph is defined by a set of nodes or vertices *V*, and an edge set *E*. In the context of this application, the set *V* = {***v***_***i***_|*i* = 1, … *N*} consists of set of patches contained in the whole slide images. Each node ***v***_***i***_ = (***g***_***i***_, ***h***_***i***_) encodes the spatial location (***g***_***i***_) and feature representation (***h***_***i***_) of a patch, where ***h***_***i***_ ∈ **ℛ**^**1024**^ is 1024-dimensional feature representation of a patch extracted from ShuffleNet [50] pretrained on ImageNet [51]. The edge set *E* is obtained by linking each node with their neighbouring nodes (within 4000 pixels) using Delaunay triangulation. An edge *e*_*ij*_ ∈ *E* exists in the set *E* if two nodes *v*_*i*_ and *v*_*j*_ are connected.

## 3.4 Prediction of drug sensitivity using Graph Neural network (GNN)

We utilize a Graph Neural Network (GNN) to predict node-level and WSI-level sensitivity of patients to a set of drugs *D* from their corresponding WSI-graphs. More specifically, we used

*SlideGraph*^*∞*^with slightly modified architecture [37]. The multi-output GNN model predicts patch-level and WSI-level sensitivity of patients to *D* different drugs in an end-to-end manner. Node-level representation (feature embedding of a patch) undergoes a series of EdgeConv layers *L* = {1,2,3}. In each EdgeConv layer [52], the representation of each node in the graph is updated by gathering information from its neighbouring nodes using message passing. This aggregation process generates embeddings that are utilized in the subsequent layers. The output embedding of an EdgeConv for a node at index *m* in layer *l* can be mathematically written as follows:

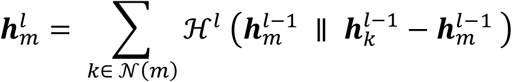

In the above equation, the term 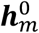 is the initial embedding of node *m* which is equivalent to ***h***_*m*_. The symbol 𝒩(*m*) represents the neighbouring nodes of *m* and ℋ^*l*^ denotes a multi-layer perceptron at layer *l*. The EdgeConv operation updates the feature representation of a node 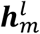 by aggregating information from its neighbouring nodes 𝒩(*m*). In the case of |*L*| = 3, each node is expected to gather information from the neighbouring nodes that are 4-hops apart in the WSI-level graph.

The node-level embedding 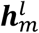 of a node ***v***_*m*_ = (***g***_*j*_, ***h***_*j*_) ∈ *V* is passed as input to a multi-layer perceptron 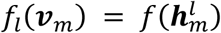 for generating node-level prediction score. The patch-level prediction score of patient sensitivity to different drugs can be obtained by aggregating node-level prediction score across all layers 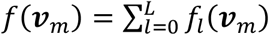. WSI-level prediction score *F*(*G*) is then obtained by pooling and aggregating node-level prediction score *F*(*G*) = ∑_∀ *m*∈*V*_ *f*(***v***_*m*_).

The trainable parameters of both EdgeConv layers and node-level regressor are learned in an end-to-end manner using backpropagation. During training, for a batch of size *N* = 16 patients, the predicted sensitivity values for *d* = {1 … *D*} drugs are compared with ground-truth values using pairwise ranking loss, mathematically formulated as follows:

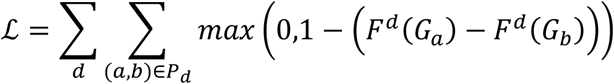

In the above formulation 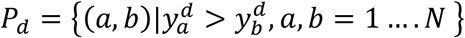 represents all pairs (*a,b*) where the ground-truth sensitivity estimate of patient *a* is greater than patient *b* for drug *d*. Minimizing the loss function will enforce the model to rank highly sensitive patients higher than the low sensitive ones for all drugs.

## 3.5 Model training and evaluation

We trained and evaluated the performance of the proposed method using 5-fold cross-validation in which the data was partitioned into non-overlapping 80/20 training and test splits. For validation, we randomly selected 20% data from the training set and use it for parameter tuning and optimization. We trained the model for 300 epochs using adaptive momentum-based optimizer [53] with a learning rate of 0.001 and weight decay of 1e-4 on the training set using a batch size of 16. We stopped the model training if the validation loss is not minimizing over 20-consective epochs. During training, we used a queue of size 10 and put the best model in the queue based on its performance over the validation set. For the test set, we aggregated the scores of the 10-best models from the queue to obtain a final prediction score. To evaluate the model performance for a given drug on the test set, we computed Spearman Correlation Coefficient (SCC) value between the ground-truth and predicted sensitivity values with its associated p-value. For a given drug, the *p*-value associated with SSC values across multiple cross-validation runs were combined by calculating twice the median p-value (p50) as a conservative estimate for statistical significance [54]. For predictive performance evaluation, we used the *p*-value and mean SCC as performance metrics.

## 3.6 Identification of histological patterns associated with drug sensitivity

To identify histological patterns associated with high and low sensitivity of a certain drug, we divided patients into two classes (high sensitivity and low sensitivity). For each class, we selected top 50 patients based on absolute difference between imputed sensitivities and model predicted sensitivities. From the WSIs of patients belonging to highly sensitive groups we extracted the highest scoring (based on node-level score) 1% patches, while for low sensitive cases we extract the lowest scoring 1% patches. Within each class, we then clustered the patches to uncover visual patterns associated with high and low sensitivity using 25-medoid clustering [55]. After clustering, we obtained 25 visual patterns representative of high and low sensitivity of a certain drug.

## 3.7 Cellular composition and statistical analysis

We analysed the cellular composition of high and low scoring patches in their respective high and low sensitive group using our in-house state of the art cellular composition predictor ALBRT [29]. For a given patch, ALBRT generated a 4-dimentional vector representing the counts of neoplastic, inflammatory, connective, and epithelial cells present in a patch. ALBRT was originally trained on patches of size 256 × 256 pixels at a spatial resolution of 0.25 MPP, so we tiled each patch of size 512 × 512 pixels into 4-subpatches and aggregated the ALBRT predicted cellular composition. The patch-level inflammatory to neoplastic cell ratio was computed based on the cellular composition. This was done by dividing the count of inflammatory cells by the sum of neoplastic and inflammatory cell counts.

## 3.8 Estimation of mitotic counts

The mitosis detection was done using the state-of-the-art mitosis detection method called MDFS (mitosis detection: fast and slow) [56]. The MDFS method follows a two-stage approach to detect mitotic candidates. It first detects the mitotic candidates using a convolutional neural network (CNN) and then subsequently refines the prediction by training a CNN classifier. For more details interested user is referred to MIDOG challenge paper [27]. After detecting the mitotic figures, we estimated the patch-level mitotic counts by counting all the detected mitoses in the patch.

## Data and code availability

Whole slides images (WSIs) of all TCGA-BRCA patients used in the study can be downloaded from the NIH Genomic Data Common Portal at this link: https://portal.gdc.cancer.gov/. Histological analysis was performed in python. The code of modified version of *SlideGraph*^*∞*^used in the study can be found at: https://github.com/engrodawood/Hist-DS.

